# Cross-talk between SIM2s and NFkB regulates cyclooxygenase 2 expression in breast cancer

**DOI:** 10.1101/634113

**Authors:** Garhett Wyatt, Chloe Young, Lyndsey Crump, Veronica Wessells, Tanya Gustafson, Yang-Yi Fan, Robert Chapkin, Weston Porter, Traci R Lyons

## Abstract

**Background:** Breast cancer is a leading cause of cancer-related death for women in the United States. Thus, there a need to investigate novel prognostic markers and therapeutic strategies. Inflammation raises challenges to both treating and preventing the spread of breast cancer. Specifically, the nuclear factor kappa b (NFkB) pathway contributes to cancer progression by stimulating proliferation and preventing apoptosis. One target gene of this pathway is *PTGS2*, the gene that encodes for cyclooxygenase 2 (COX-2), which is upregulated in 40% of human breast carcinomas. COX-2 is an enzyme involved in inflammation. Here we investigate the effect of Singleminded 2s, a transcriptional tumor suppressor that is implicated in inhibition of tumor growth and metastasis, in regulating NFkB and COX-2.

**Methods:** We utilized in vitro reporter assays, immunoblot analyses, qPCR and immunohistochemical analysis to dissect the relationship between NFκB, SIM2s, and COX-2. Furthermore, we utilized COX-2 targeting strategies to determine tumor suppressive activities.

**Results:** Our results reveal that SIM2s attenuates the activation of a NFκB via luciferase reporter assays. Furthermore, immunostaining of lysates from breast cancer cells over expressing SIM2s showed reduction in various NFκB signaling proteins, whereas knockdown of SIM2 revealed increases in the same NFκB signaling proteins. Additionally, by increasing NFκB translocation to the nucleus in DCIS.COM cells, we show that NFκB signaling can act in a reciprocal manner to decrease expression of *SIM2s*. Likewise, suppressing NFκB translocation in DCIS.COM cells increases *SIM2s expression*. We also found that NFκB/p65 represses *SIM2* in via dose-dependent manner and when NFκB is suppressed the effect on the *SIM2* is negated. Additionally, our CHIP analysis confirms that NFκB/p65 binds directly to SIM2 promoter site and that the NFκB sites in the SIM2 promoter are required for NFkB-mediated suppression of SIM2s. Finally, over expression of SIM2s decreases *PTGS2* in vitro and COX-2 staining *in vivo* while decreasing *PTGS2* and/or Cox-2 activity results in re-expression of SIM2. Our findings identify a novel role for SIM2s in NFκB signaling and COX-2 expression.

**Conclusions:** These findings provide evidence for a mechanism where SIM2s may represses COX-2 expression to provide an overall better prognosis for breast cancer patients.

## Background

Despite improvement in early detection and treatment, breast cancer remains the second leading cause of cancer-related death for women in the United States [1]. Breast ductal carcinoma in situ, referred to as DCIS, consists of a heterogeneous group of diseases characterized by a neoplastic mammary lesion that is confined to the ductal system of the breast [2]. DCIS progresses to invasive ductal carcinoma (IDC) through events such as epithelial mesenchymal transition (EMT), basement membrane degradation, controlled inflammatory signaling and other pathways associated with a wound-healing milieu [3–5]. It is estimated that ~20% of mammography detected breast cancers are DCIS[6] and ACS Facts and Figures indicates ~65,000 cases of DCIS are diagnosed per year[7]. Provided that DCIS is removed surgically, as is standard of care, a woman diagnosed with DCIS without recurrence is more likely to die of other causes than of breast cancer[8]. However, it is estimated that ~15-20%, or ~10-13,000, DCIS patients recur with invasive disease within a decade[9, 10]. Recently identified risk factors for DCIS recurrence include age<40 at diagnosis, African American ethnicity, hormone receptor negativity, and HER2 positivity[8]. However, these high-risk groups only account for 20% of the DCIS patient population[10]. Therefore, identifying additional risk factors for, or markers that will predict, DCIS aggressiveness are extremely important goals for preventing invasive cancer in DCIS patients.

There is increasing evidence that inflammation plays a key role in breast cancer progression [11]. One such specific inflammatory pathway is nuclear factor kappa b (NFκB). The NFκB signaling pathway includes five members: NFKB1 (p105/p50), NFKB2 (p100/p52), RelA (p65), RelB and c-Rel. Dimers of the aforementioned proteins are held in the cytoplasm by inhibitor kappaB (IkB) proteins, primarily the IkBa protein. The mechanism of action for NFκB includes phosphorylation of IkBa by inhibitor of kappaB kinase (most commonly IKKα and IKKβ), which leads to degradation of IkBa. Upon degradation of the IkBa, NFκB heterodimers, specifically the canonical heterodimer p50/p65, translocate to the nucleus and bind to promoters of certain target genes, leading to activation of transcription[12, 13]. Two NFκB consensus sites are located in the promoter region of the human *PTGS2* gene, which encodes for pro-inflammatory enzyme COX-2[14]. These NFκB sites are conserved on the 5’-promoter regions in both mouse and human, however the 3’ promoter is only found in human[15]. These NFκB consensus sites not only contribute to cancer progression by stimulating and preventing apoptosis, but also the activation of cyclooxygenase 2 (COX-2) mediated signaling [16]. COX-2 is the inducible form of cyclooxygenase, which is the key enzyme involved in the biosynthesis of the pro-inflammatory agent prostaglandin[17–22]. COX-2 has been implicated in DCIS progression through promotion of mammary tumorigenesis via increases in proliferation, migration, invasion and metastatic spread in pre-clinical models[23–25]. Additionally, expression of COX-2 is frequently observed in patients with invasive disease and is associated with DCIS recurrence. Furthermore, therapeutic benefit of inhibiting COX-2 has been observed in colon, esophagus, lung, bladder, breast and prostate cancers[19, 20, 26–36]. Thus, it is logical to expect that inhibition of COX-2 in breast cancer patients could enhance overall prognosis.

We have shown that Singleminded-2s (SIM2s; expressed from *SIM2*), a member of the bHLH/PAS family of transcription factors, is a tumor suppressor that is expressed in breast epithelial cells and down-regulated in the transition from DCIS to IDC [37–40]. Specifically, using the DCIS.COM progression model we demonstrated that re-expression of *SIM2s* inhibits growth, invasive phenotypes, and progression to metastasis. Furthermore, expression of SIM2s promotes a more luminal-like phenotype in breast cancer cells whereas down-regulation of *SIM2s* leads to an increase in invasive potential [40]. Consistent with the role for SIM2s in cancer progression, we have also shown that the NFκB signaling pathway is downregulated by SIM2s over expression during mammary development, specifically during postpartum mammary involution[41]. In this study, we expand on these observations and demonstrate a relationship between SIM2s, the NFκB signaling pathway, and COX-2. We suggest that re-expression of SIM2s could be mediated by inhibition of COX-2 signaling, which may serve to reduce breast cancer progression.

## Methods

### Cell Culture

MCF7 and Sum159 cells were purchased from American Type Culture Collection (ATCC) and were maintained in accordance to their guidelines. MCF10-DCIS.COM cells were generously donated by Dr. Dan Medina (Baylor College of Medicine, Houston, TX, USA). Cells were plated in 6 well plates for RNA isolation experiments according the guidelines from ThermoFisher Scientific. Celecoxib experiment were performed as follows; cells were first plated in 10uM celecoxib for 24hours, then media was changed and treatment was performed at 20uM celecoxib for 24hours and then harvested for analysis. DHA experiments on cell lines were performed as follows; cells were dosed with 50uM DHA for 24 hours and then harvested for analysis.

### Generation of cell lines

Point mutations in the SIM2 gene were generated via long cDNA synthesis (Invitrogen). Plasmids were amplified using Subcloning Efficiency DH5a competent cells (Life Technologies). Plasmid DNA was isolated using the HiPure Plasmid Maxiprep kit (Life Technologies) or the ZymoPURE Plasmid DNA Isolation Kit (Zymo Research). Viral transduction was then performed as previously described [39]. Puromycin selection (2ug/mL) was started the following day and maintained for a week.

### Animal Models

200,000 MCF10DCIS cells stably expressing anti COX-2 shRNAs (a generous gift from Kornelia Polyak and Andriy Marusyk) were orthotopically injected and tumors harvested as previously described [23, 24].

### RNA isolation and real-time PCR

As previously described [42]. Primers can be found in Supplementary Table 2.

### Immunoblotting

Cells were washed with cold PBS and lysed in high-salt lysis buffer (50mM HEPES, 500mM NaCl, 1.5mM MgCl2, 1mM thylenediaminetetraacetic acid (EDTA), 10% glycerol, 1% Triton X-100, pH 7.5) supplemented with 1 mM Na_3_VO_4_ (Sigma) and 1mM complete ULTRA tablets mini EDTA-free Easy pack (Roche). Protein concentration was determined using the DC Protein Assay (Bio-Rad) with bovine serum albumin as a standard. Immunoblotting and zymography were then conducted as previously described[39]. Antibodies can be found in supplementary Table 1. Blots were imaged on a ChemiDoc MP (Bio-Rad) after incubating in ProSignal Pico ECL Spray (Genesee Scientific) for 3 minutes. Quantification performed using ImageJ.

### Immunohistochemistry

IHC for COX-2 was performed as previously described [23]. Analysis for positive nuclei was performed previously described[25]. Antibodies can be found in supplementary Table 1.

### Transient transfection

293T cells were used for all transfections for luciferase activity. One hundred ng (0.1 μg) of plasmid containing transcription factor was mixed with up to 1μg (amount varies per construct) of plasmid containing promoter construct. Three μl of Genejuice (Novagen) was used per microgram of DNA. DNA and Genejuice were mixed in Opti-MEM media (Invitrogen). Protein was harvested 2 days after transfection, using Reporter Lysis Buffer (Promega). Luciferase activity and total protein were measured as described previously [38]. Luciferase activities were normalized to total protein values and are represented as the means ± SE for three wells per condition.

### Chromatin Immunoprecipitation

For ChIP assays, 2μg of plasmid containing repressor and 2μg of plasmid containing the SIM2 promoter construct were transfected into 293T cells on a 10cm plate. Chromatin was harvested 2 days after transfection.

### Statistical Analysis

To address scientific rigor, all experiments in cell lines and xenografts were conducted in biological triplicates at a minimum, technical duplicates, and repeated three times. Normal distribution was confirmed before conducting unpaired *t* test. Significance was considered at *p* <0.05 unless otherwise noted.

## Results

### SIM2s downregulates NFKB signaling

To test the hypothesis that SIM2 directly affects NFκB/p65 mediated transcription, we co-transfected a reporter plasmid encoding a NFkB binding site upstream of the luciferase gene (5X NFκB-luc) with the p65 subunit along with SIM2s in 293T cells and measured relative light units as a readout for NFκB activity. As expected, p65 strongly activated the reporter construct; however, this response was blocked by co-transfection of *SIM2s* (Fig 1A). To determine the mechanism of this inhibition, the transfection was repeated with a SIM2s expression construct missing the Pro/Ala transcriptional repression domain (SIM2sΔR). This construct also significantly attenuated the activation of the 5X NFκB-luc construct by NFκB/p65, demonstrating that the repression domain of SIM2s is not required for inhibition of NFκB signaling (Fig 1B). Alternatively, to determine whether SIM2 modulates expression levels of mediators of the NFκB pathway in our breast cancer cell lines to downregulate signaling, we performed western blot analysis and found that IKKa, IKKb, phosphorylated-p65 and p65 protein levels were all decreased in *SIM2s* overexpressing SUM159-cells (Figure 1C). Similarly, we found that NFκB pathway protein levels were increased in SIM2s knockdown MCF7 cells (Figure 1C). These results suggest that SIM2s may affect NFκB mediated transcription via modulation of expression of key mediators of NFκB signaling.

**Figure 1.**
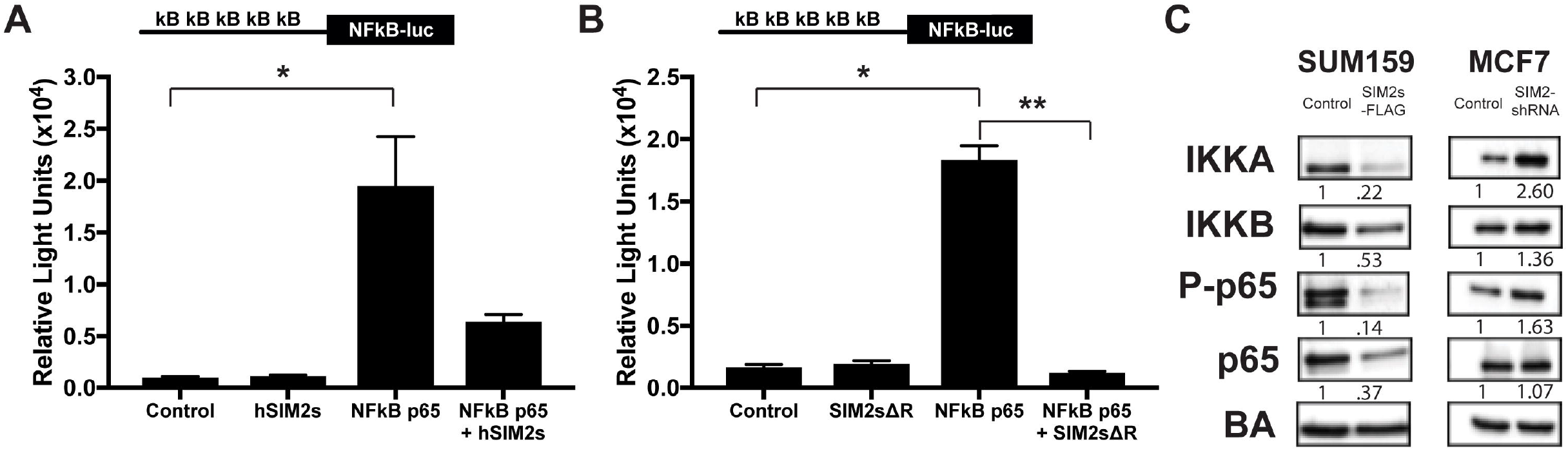
**A.** Luciferase activity in HEK293T cells co-transfected with 5x kB binding sites upstream of the luciferase gene (5x NFkB-luc) and NFkB p65 and/or SIM2s. (Diagram of promoter construct is shown above for reference.) **B.** Luciferase activity in HEK293T cells co-transfected with 5x NFkB-luc and NFkB p65 and/or SIM2s with its repression domain deleted (SIM2sΔR). **C.** SUM159 plpcx emp (control), SUM159 plpcx SIM2s-FLAG (overexpression), MCF7 psil SCR (control), and MCF7 psil SIM2-shRNA (knock down) were analyzed by immunoblot for levels of IKKA, IKKB, phospho-p65, p65 and Beta Actin as loading control. Unpaired *t* test: *, *P* <0.02; **, *P*<0.05. Analysis performed via ImageJ for comparison of protein expression.

### NFκB signaling downregulates SIM2s expression

We also observed that stable over expression of an inhibitor of the inhibitor kappa kinase beta (IKKβ), which normally retains NFκB in the cytoplasm, significantly decreases SIM2s gene expression in the DCIS.COM cells (Figure 2A). Likewise, when the NFκB pathway is suppressed via stable transduction of the inhibitor of kappaB alpha super repressor (IκB-SR), which maintains the NFκB heterodimer (p50/p65) in the cytosol, *SIM2s* expression was increased (Figure 2B). To test the hypothesis that NFκB downregulates *SIM2* expression, we cloned a 2kb portion of the *SIM2* promoter upstream of the luciferase gene and co-transfected with increasing amounts of p65 in HEK293T cells. We observed dose-dependent repression of SIM2s promoter activity (Figure 2C). Importantly, co-transfection with IκB-Super Repressor (IκB-SR), as well as IκB-SR with NFκB p65, successfully reversed the downregulation of SIM2s promoter activity (Figure 2D) suggesting that this was not a dominant negative effect. Subsequently, analysis of the *SIM2* promoter identified two consensus NFkB binding sites near the transcriptional start site for *SIM2*. Thus, we utilized chromatin immunoprecipitation (ChIP) analysis to show that p65 directly binds to the SIM2 promoter around the transcriptional start site (Figure 2E). Then, to determine whether binding of p65 to the NFkB binding sites is necessary for downregulation of SIM2s expression, we mutated the two NFκB sites in the *SIM2* promoter reporter construct and performed additional co-transfection experiments with p65. Interestingly, the NFκB double mutant promoter failed reduce SIM2 promoter activity when compared to the wild-type promoter (Figure 2F), implicating a direct interaction of NFκB/p65 to decrease SIM2 transcription. These results suggest that NFκB mediated transcriptional repression of SIM2s may allow for activation of NFκB mediated transcription of pro-inflammatory target genes such as *PTGS2*.

**Figure 2.**
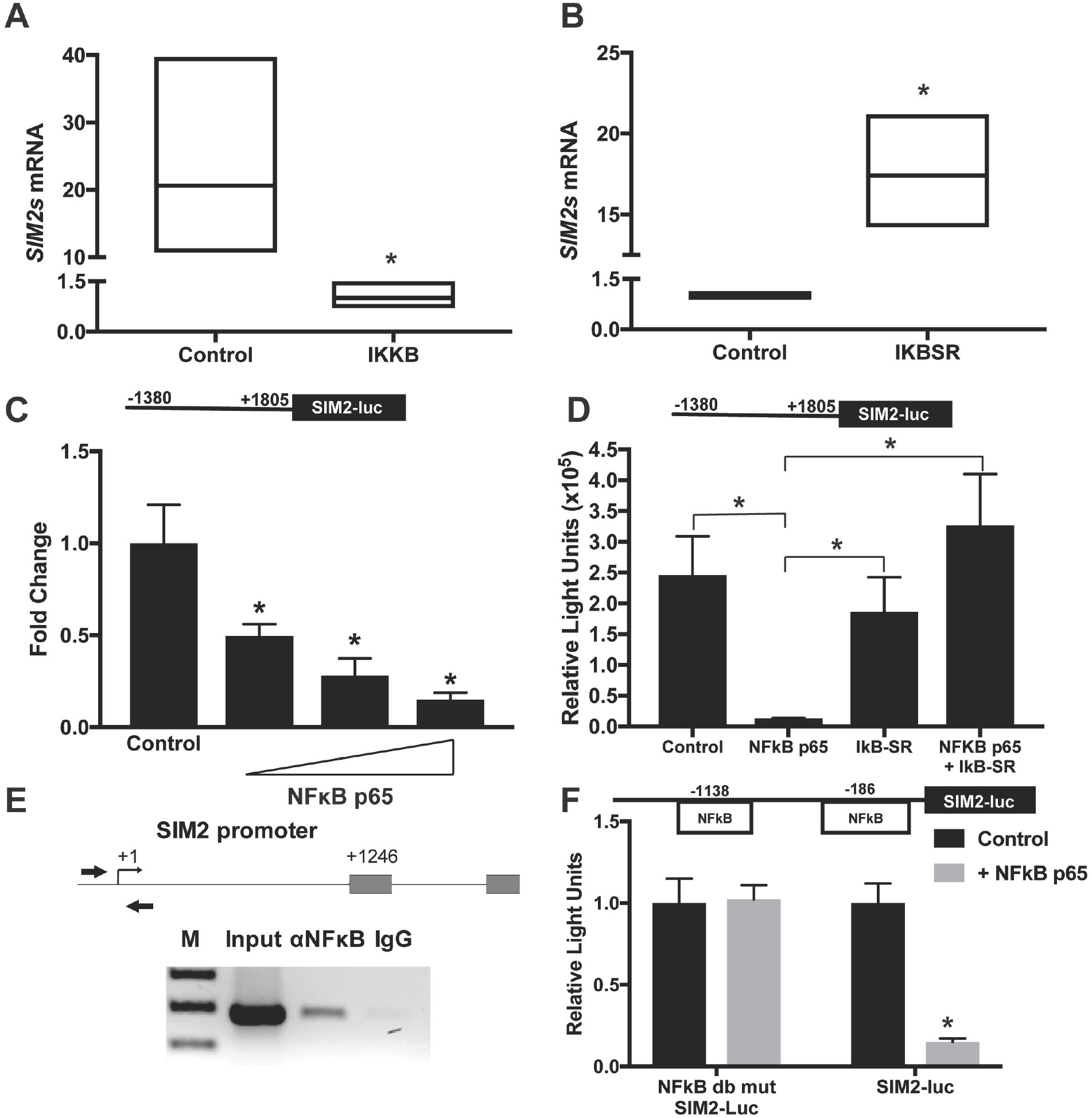
**A.** *SIM2s* expression in DCIS.COM control cells and cells overexpressing IKKB by qPCR as fold change. **B.** *SIM2s* expression in DCIS.COM control cells and cells overexpressing IKB-SR by qPCR as fold change. **C.** SIM2 promoter activity in HEK293T cells co-transfected with SIM2 promoter upstream of the luciferase gene and increasing amounts of NFκB p65 (50uM,100uM, and 150uM). **D.** SIM2 promoter activity after co-transfection of promoter with control vector (pcDNA3), NFκB p65, and/or IκB-SR. **E.** ChIP assay for NFκB p65 binding after transient transfection of SIM2 promoter with NFκB p65 in HEK293T cells. **F.** SiM2 promoter activity in HEK293T Cells co-transfected with SIM2 promoter upstream of the luciferase gene and 150uM NFκB p65 compared with the SIM2 promoter activity in HEK293T cells co-transfected with NFκB double mutant SIM2 promoter upstream of the luciferase gene. Unpaired *t* test: *, *P*<0.05.

### SIM2s expression downregulates COX2

To explore the relationship between SIM2s and *PTGS2* expression in breast cancer we analyzed three different breast cancer cell lines including MCF7, MCF10ADCIS.COM (DCIS.COM), and SUM159 cells. The non-invasive MCF7 cell line and the highly invasive triple-negative SUM159 cell line were utilized to examine the differential expression of SIM2, and subsequent regulation *PTGS2*, as it relates to invasion. DCIS.COM cells were also used for their unique ability to mimic basal-like DCIS *in vivo* and their ability to progress to invasive disease upon acquisition of COX-2 protein expression[23, 43]. We have previously shown that the invasive competent DCIS.COM cells have more *SIM2* expression when compared with the non-invasive MCF7 [38, 39]. Confirming and extending this observation, qPCR analysis reveals lowest *PTGS2* expression in MCF7 cells, which was increased 130-fold in DCIS.COM cells and highest in in the SUM159 cells (SFigure1A). To determine whether reduction of *SIM2s* in the non-invasive cells could increase expression of *PTGS2*, we analyzed control and *shRNA-SIM2s* DCIS.COM and MCF7 cells by qPCR. Our results revealed that downregulation of *SIM2s* led to a significant increase in *PTGS2* gene expression in both cell lines (Fig 3A, Fig 3B). Moreover, we found that over expression of *SIM2s* in highly invasive SUM159 cells significantly inhibited *PTGS2* expression (Fig 3C). In our previous studies, we showed that over expression of *SIM2s* in DCIS.COM cells blocked invasion in vivo, whereas loss of *SIM2s* or over expression of the protein product of *PTGS2*, COX-2, resulted in increased invasion and metastasis[23, 42]. To determine the relationship between SIM2s and COX-2 protein expression, we performed immunohistochemical (IHC) analysis for COX-2 in tumors derived from control and *SIM2s* DCIS.COM xenografts to show that COX-2 levels were decreased with over expression of *SIM2s* in vivo (Fig 3D). Taken together these results suggest that SIM2s may repress invasion in the DCIS.COM model by promoting downregulation of COX-2.

**Figure 3.**
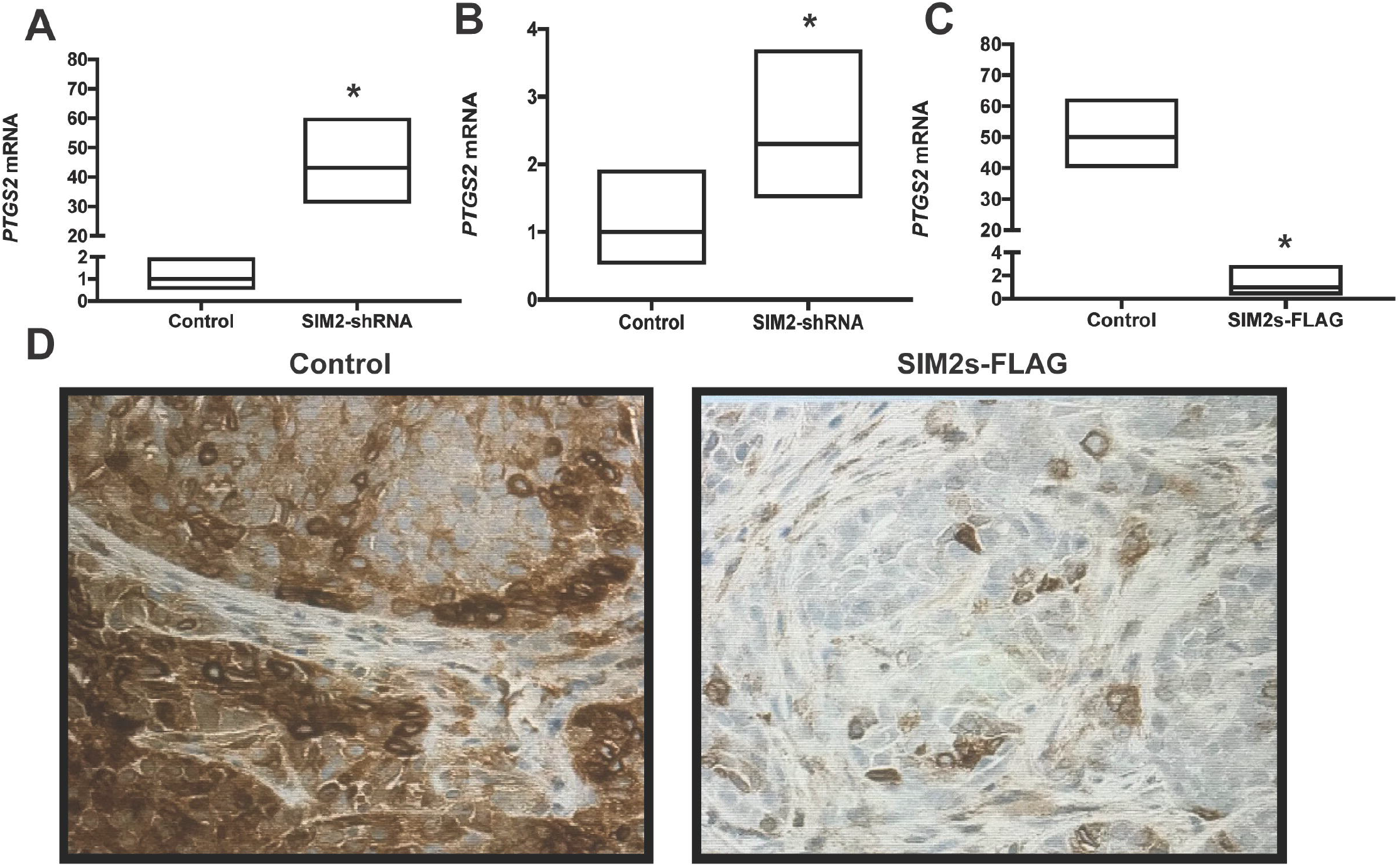
**A.** *PTGS2* expression in MCF7 control cells and cells overexpressing SIM2s by real time qPCR as fold change. **B.** *PTGS2* expression in DCIS.COM control cells and cells with SIM2-shRNA by real time qPCR as fold change. **C.** *PTGS2* expression in SUM159 control cells and cells overexpressing SIM2s by real time qPCR as fold change. **D.** Immunohistochemistry for COX-2 in DCIS.COM cells stably transduced with control vector, SIM2s-FLAG (overexpression), respectively. Unpaired *t* test: *, *P* <0.01.

### COX-2 downregulation restores SIM2s

Since the invasive potential in DCIS.COM positively correlates with, and depends upon expression and activity of, COX-2 [23] we proposed to test the hypothesis that the loss of invasive phenotype observed with blocking of COX-2 expression and/or activity was due, in part, to re-expression of *SIM2s*. Thus, we measured SIM2 protein levels in cells with confirmed stable knockdown of PTGS2/COX-2 via immunoblot. Indeed, DCIS.COM *shPTGS2* and control cells exhibited and inverse relationship between SIM2s and COX-2(Figure 4A). Extending these observations IHC analysis of tumors generated from control and *shPTGS2* DCIS.COM cells, which are less invasive[23], revealed an increase in positive nuclear staining for SIM2 with *PTGS2* knockdown (Figure 4B&C). To determine whether COX-2 activity drives the inverse relationship between *SIM2* and *COX-2* and cell invasion, we treated the highly invasive SUM159 cells with a dose of the selective COX-2 inhibitor, celecoxib, that had previously been shown to decrease invasion of COX-2 expressing cells[23]. We observed a significant increase in SIM2 expression (Figure 4D). Additionally, we show that Docosahexaenoic (DHA), a n-3 polyunsaturated fatty acids (PUFA) that can result in a shift to a more anti-inflammatory gene expression profile[44], and can reduce COX-2 expression[45–48], significantly increases *SIM2s* expression (Figure 4E). Thus, our driving hypothesis is that reduction of inflammatory pathways via inhibition of activity and/or decreased COX-2 expression results in re-expression of *SIM2s* and may be one mechanism for preventing progression of DCIS to invasive breast cancer[23].

**Figure 4.**
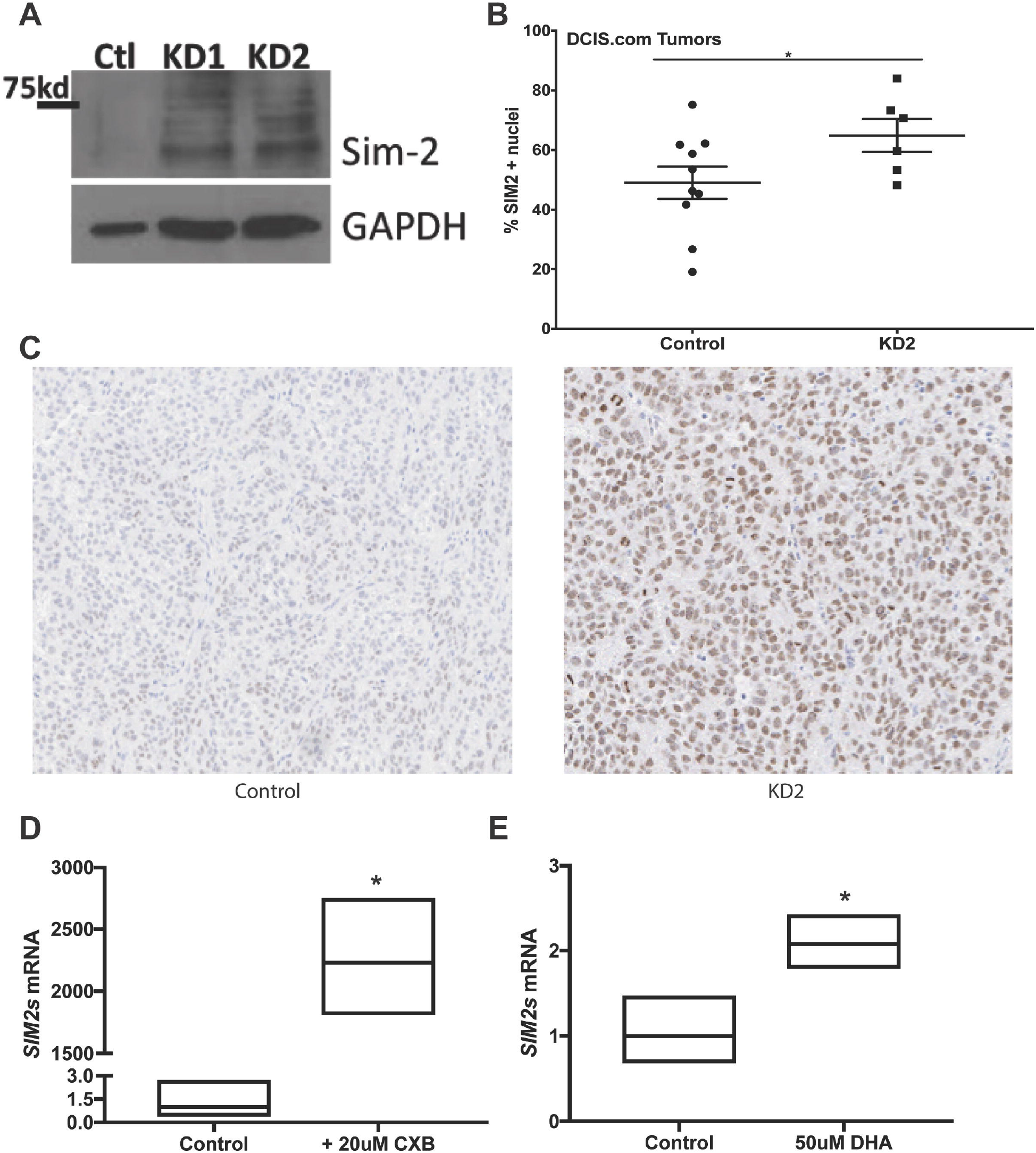
**A.** DCIS.COM control, 1 *shPTGS2*, 2 *shPTGS2* were analyzed by western blot for SIM2 and GAPDH as loading control. **B.** IHC Analysis for SIM2 positive nuclei in tumors generated from control and *shPTGS2* DCIS.COM cells. **C.** Images of IHC analysis for SIM2 in Tumors generated from control and *shPTGS2* DCIS.COM cells. **D.** *SIM2s* expression in SUM159 control cells and cells dosed with 20uM Celecoxib by qPCR as fold change. **E.** *SIM2s* expression in DCIS.COM control cells and cells dosed with 50uM DHA by qPCR as fold change. Unpaired *t* test: *, *P*<0.05.

## Discussion

Through transgenic mouse models and in vitro studies SIM2s has been identified as a novel player in several key aspects of mammary gland development. Specifically, genetic ablation of SIM2s in mammary epithelial cells revealed that SIM2s is required for ductal morphogenesis and differentiation of luminal cells for milk production during lactation. Furthermore, mammary specific overexpression of SIM2s resulted in a delay in post lactational mammary gland involution through suppression of Stat3 and NFKB signaling, as well as maintenance of a markers of epithelial cell differentiation normally observed only during lactation. These results suggest that SIM2s has tumor suppressive activities in the mammary gland through maintenance of epithelial cell differentiation. Consistent with this, loss of *SIM2s* expression in the mammary epithelium results in epithelial to mesenchymal transition (EMT) events, such as loss of E-cadherin and increases in matrix metalloprotease activity; results which are also observed in breast cancer cell lines. SIM2s is also downregulated in breast cancer patient samples further validating its potential role in tumor suppression[42]. In this study, we demonstrate a novel role for SIM2s as a negative regulator of the NFκB pathway, which normally results in downstream transcriptional activation of COX-2 expression. We have also shown that the SIM2s is targeted for suppression by NFκB signaling suggesting a regulatory feedback loop. Of particular interest, loss of *SIM2s* drastically increases pro-tumorigenic COX-2 expression and loss of COX-2 activity and expression results in re-expression of SIM2s. Thus, we have identified a reciprocal relationship between a molecule with known tumor suppressive activities, SIM2s, and known tumor promotional molecules NFκB and COX-2(Figure 5).

**Figure 5.**
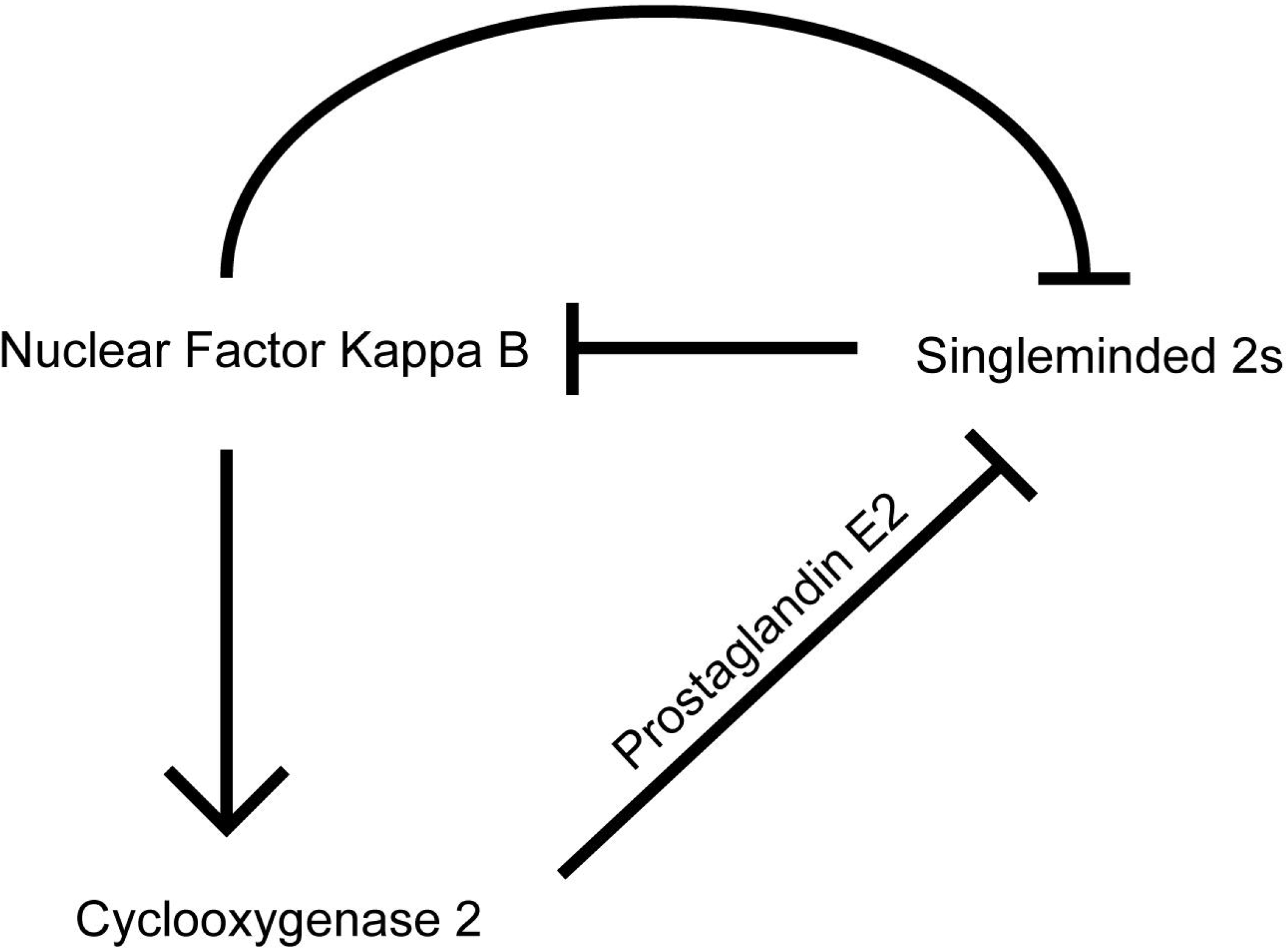
A model depicting NFkB-mediated SIM2 signaling. Our current data suggest that NFκB inhibits SIM2 expression through an unknown mechanism to result in upregulation of COX-2 expression and subsequent activity, which further downregulates SIM2. SIM2 is also capable of decreasing NFκB signaling to down regulate COX-2 expression.

We predict that loss of SIM2s may be important for progression of in situ lesions to invasive disease. In a model of DCIS, loss of *SIM2s* is associated with an increase in invasiveness, enhanced tumor aggressiveness and progression. Specifically, upon loss of SIM2 in tumors an increase in colocalization of keratin 5 and vimentin have been observed[42], which is indicative of mesenchymal and invasive phenotypes. Additionally, matrix metalloproteinases (mMPs), which are associated with basement membrane degradation during mammary gland development and cancer, are also significantly increased with loss of SIM2[49–51]. Furthermore, P21, an important cell cycle regulator involved in both the p53 stress-response pathway and senescence, is markedly decreased mRNA in SIM2 knockdown xenografts[52, 53]. These changes likely promote an increased potential for progression of DCIS to IDC. Furthermore, increased COX-2 coupled with increased p16 and increased proliferation are associated subsequent recurrence of DCIS[22]. Here we show that cells with low invasive potential, the DCIS.COM and MCF7 cells, exhibit increased expression of COX-2 upon knockdown of *SIM2s* and endogenously express moderate levels of SIM2 compared to the low level of SIM2 observed in the more invasive SUM159 cells[38]. Likewise, overexpression of SIM2s in SUM159 cells decreases COX-2 expression. Coincidently, in a DCIS.COM xenograft study, SIM2s overexpression also significantly decreased COX-2 staining in tumor sections and xenografts from MCF10DCIS shCOX-2 cells exhibit increased expression of SIM2. These data demonstrate, for the first-time, a link between SIM2s expression and COX-2 and provide evidence of negative feedback of COX-2 on SIM2 expression. Since it is well known in the literature that COX-2 inhibition is associated with better prognosis for breast cancer patients[54, 55], further studies on strategies for re-expression of SIM2s may be beneficial in improving prognosis of breast cancer patients. Furthermore, an additional implication is that SIM2s could be utilized as amarker to identify DCIS patients that are of low risk for acquisition of COX-2 expression and progression to IDC.

## Conclusions

These findings support a role for SIM2s in the prevention of breast cancer progression through its ability to interact with *PTGS2* expression via modulating the NFκB signaling pathway. It has long been established that NFκB regulates genes involved in cell proliferation and cell survival. Specifically, blocking NFκB in tumor cells can lead to susceptibility from anti-cancer agents. However, due to the complexity of the tumor micro environment, NFκB signaling also has been found to have anti-cancer effects in various cancer cells. Thus, it is important to investigate a mechanism, specifically in mammary tissue, in which the targeted pathways are highly involved with cell proliferation, survival, migration and invasion. Due to elevated COX-2 expression correlating with poor prognosis, it is imperative to investigate reducing COX-2/*PTGS2* expression. In the data provided here, we have demonstrated an integral role for SIM2s involvement in mediating NFκB signaling to decrease expression of COX-2/*PTGS2* leading to an improved prognosis for breast cancer patients.

## Abbreviations

CHIP: Chromatin immunoprecipitation
COX-2: Cyclooxygenase 2
DCIS: Ductal carcinoma in situ
DHA: Docosahexaenoic acid
EMT: Epithelial mesenchymal transition
IDC: Invasive ductal carcinoma
IκB: Inhibitor kappa b
IκBα: Inhibitor kappa b alpha
IκB-SR: Inhibitor kappa b alpha super repressor
IKKα: Inhibitor kappa b kinase alpha
IKKβ: Inhibitor kappa b kinase beta
NFκB: Nuclear factor kappa b
PUFA: polyunsaturated fatty acids
SIM2s: Singleminded-2s

## Declarations

## Acknowledgements

We gratefully acknowledge Kornelia Polyak and Andriy Marusyk for providing the shCOX-2 DCIS cell lines

## Availability of data and materials

Not applicable

## Authors’ contributions

GW performed the in vitro experiments, formal data analysis, interpretation of the data, production of figures, writing of the original draft, as well as reviewing and editing the final draft of the manuscript.CY and LC were involved in tumor immunostaining and immunoblot analysis. VW was involved in designing and performing immunostaining of the xenograft studies. TG was involved in animal husbandry, in vivo experiments and CHiP analysis. YF and RC provided the polyunsaturated fatty acid resources. WP and TL oversaw all the experiments and experimental design, interpretation of the data, provided the resources, and contributed to the writing as well as reviewing and editing of the manuscript. All authors read and approved the final manuscript.

## Ethics approval and consent to participate

Animal studies were approved by the Institutions laboratory care committee.

## Consent for publication

Not applicable

## Supplemental Figure Legends

**SFigure 1.** qRTPCR analysis of PTGS2 expression in 3 different breast cancer cell lines.

**STable 1.** Antibody list for immunoblot and immunohistochemical analyses.

**Stable 2.** Primer sequences for qPCR analyses.

