## Supplemental Table 2 for "Cross-talk between SIM2s and NFkB regulates cyclooxygenase 2 expression in breast cancer"

|  |  | Forward | Reverse |
| --- | --- | --- | --- |
|  | SIM2s | AGGTGGGTCAGGTCTGCTC | GAAGCAGAAAGAGGGCAAGTT |
|  | TBP | CGTCCCAGCAGGCAACA | GGTGCAGTTGTGAGAGTCTGTGA |
|  | GAPDH | CCAGGTGGTCTCCTCTGACTTC | GTGGTCGTTGAGGGCAATG |
|  | 18s | CGGCTACCACATCCAAGGAA | CTGGAATTACCGCGGCT |
|  | PTGS2 | Bio-Rad, PrimePCR | |
