## Supplemental Table 1 for "Cross-talk between SIM2s and NFkB regulates cyclooxygenase 2 expression in breast cancer"

|  | **Target** | **Manufacturer** | **Product Number** | **Dilution** | **Application** |
| --- | --- | --- | --- | --- | --- |
|  | SIM2 | Millipore | AB4145 | 1:1000 | WB |
|  | SIM2 | Aviva | Aviva ARP38551_P050 | 1:500 | WB |
|  | B-Actin | Cell Signaling Technology | 3700s | 1:5000 | WB |
|  | NFkB p65 | Cell Signaling Technology | D14E12 | 1:1000 | WB |
|  | phospho-NFkB p65 | Cell Signaling Technology | 93H1 | 1:1000 | WB |
|  | IKKβ | Cell Signaling Technology | D30C6 | 1:1000 | WB |
|  | IKKα | Cell Signaling Technology | 3G12 | 1:1000 | WB |
|  | phospho-IKKα/β | Cell Signaling Technology | 16A6 | 1:1000 | WB |
|  | phospho-Ikβα | Cell Signaling Technology | 14D4 | 1:1000 | WB |
|  | COX-2 | Cayman Chemical | 160112 | 1:400 | IHC |
|  | Anti-Rabbit Secondary | Cell Signaling Technology | 7074 | 1:2000 | WB |
|  | Anti-Mouse Secondary | Cell Signaling Technology | 7073 | 1:2000 | WB |
