## Supplementary figures and images for "Cross-talk between SIM2s and NFkB regulates cyclooxygenase 2 expression in breast cancer"

### Supplemental Figure 1

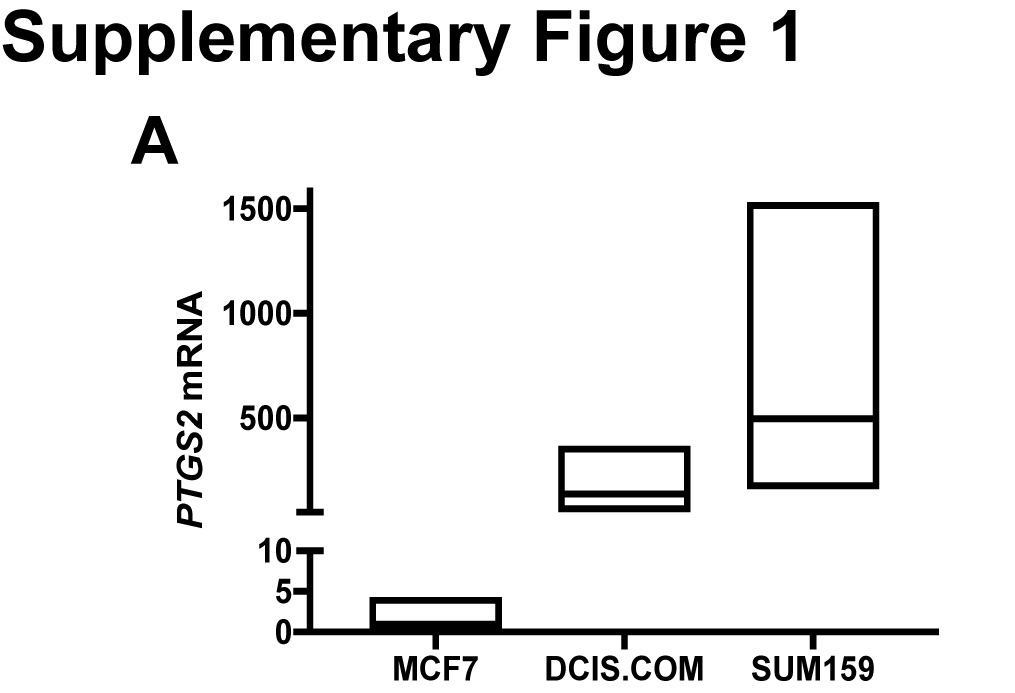
